# Genetic relateness of *Haemophilus parasuis* among reference strains and Chinese epidemic isolates

**DOI:** 10.1101/012393

**Authors:** Min Yue, Huanchun Chen

## Abstract

*Haemophilus parasuis* is the causative agent of Glässer’s disease and a commensal coloniser of the porcine upper respiratory tract. Multiple complex factors, including the early weaning of piglets and the management of high health status farms, make it a re-emerging agent, responsible for a recent increase in the prevalence and severity of disease in pigs in China. However, little genetic information is known about Chinese epidemic isolates. In this study, a phylogenetic method for genotyping the *H. parasuis* population with available Chinese epidemic isolates and reference strains from different origins is presented. Phylogenetic analysis confirmed that there are at least two different genotypes in *H. parasuis* population and a unique Chinese lineage with virulence results in the previous epidemics.

---

*Haemophilus parasuis* can colonise the porcine upper respiratory tract and also cause severe disease (Oliveira and Pijoan, 2004; Olvera et al., 2007). It is an important bacterial pathogen of pigs in China, being responsible for substantial morbidity, mortality and economic losses (Cai et al., 2005; Li et al., 2009). However, little is known about the genetic relationship between Chinese epidemic isolates, those from other countries and the reference strains. Thus, we performed a phylogenetic analysis of eighteen orthologous genes of strains of *H. parasuis* from different geographical locations and obtained from healthy and disease animals.

Eighteen orthologous genes (*gapA*, *ompP1*, *hscA*, *luxS*, *murG*, *pyrH*, *hlyX*, *ompP2*, *hemN*, *dapA*, *queF*, *lpdA*, *ksgA*, *sodA*, HAPS_1299 (conserved hypothetical protein), *smtA*, *dapB* and *ompP5*) were chosen from orthologous groups according to different functional categories. These were submitted for phylogenomic analysis, along with orthologous genes from 18 complete genomes and two draft genome sequences available for *Pasteurellaceae* (M. Yue et al., unpublished data). MEGA 4.0 was used to create Neighbour-Joining trees with interior branch test (2000 replicates; seed = 80650) (Tamura et al., 2007). The strains and their characteristics and primers used to amplify PCR products for DNA sequencing are detailed in Supplementary Tables S1 and S2, respectively.

Phylogenetically, the strains branched into two main lineages (Fig. 1), indicating that there were two independent genotypes present in the *H. parasuis* population examined. These accorded with results from analysis of *H. parasuis* population by a multilocus sequencing typing (MLST) method used by Olvera et al. (2006b); the only difference was that most of their strains in analysis originated from Europe. Genotype A contained many virulent strains, including a specific Chinese lineage associated with virulence, which also included a cluster (independent sample *t* test, *P* <0.001) containing strains that could not be typed according to the previous serological procedure (F599, F641, F603, F663, F685, F687 and F593) (Cai et al., 2005).

**Fig. 1.**
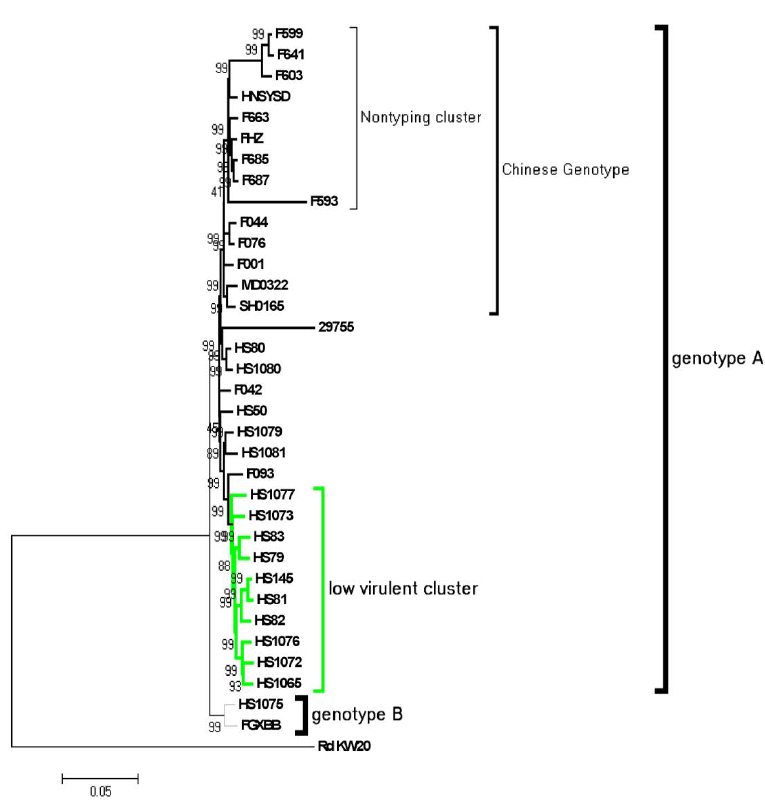
Neighbour-joining (NJ) tree with interior branch test of 2,000 replicates based on concatenated nucleotide sequences of 18 conserved genes. Thirty-four *H. parasuis* strains could be divided into two genotypes (genotypes A and B). The abundant genotype A contained the Chinese genotype, including the non-typeable cluster and a low virulence cluster (in green). Genotype B had only two strains. *H. influenzae* strain KW20 was used as the reference for the construction of the phylogenetic tree. The NJ tree was produced using MEGA 4.0. The scale bar is in unit of nucleotide substitutions per site. Bootstrap values (%) are indicated at the branch nodes of the phylogenetic trees.

Most of non-typeable strains came from central China (F641, F603, F585 and F593). The majority of Chinese epidemic isolates were composed of serovar 4, serovar 5 and non-typeable strains; the relatively high numbers of non-typeable isolates may reflect the use of serovar 4 and serovar 5 vaccines within China. The reference virulent strains (29755, HS80, HS1080, HS50, HS1079 and HS1081) and another two Chinese clinical isolates (F042 and F093) formed other virulent lineages but there was lack of convergence. However, the American Strain 29755 and HS1080 and Japanese HS80 had a closer genetic relationship with Chinese virulent lineage. The majority of epidemic strains (14/17) in the Chinese lineage (independent sample *t* test, *P* <0.001) indicated that previous *H. parasuis* in China had a genetic specificity different from other virulent lineages.

A cluster containing strains considered to be of lesser virulence (isolated from nasal cavities) formed a unique lineage (independent sample *t* test, *P* <0.001), which included avirulent strains (HS1077, HS1073, HS81 and HS1072), mid-virulent strains (HS83, HS79 and HS1065), two confirmed virulent strains (HS82 and HS1076) and an unclarified strain HS145. Interesting, the nasal isolates strains HS82 and HS1076 confirmed their virulence in animal challenge, which contradicts the knowledge that healthy nasal isolates were less virulent (Aragon et al., 2009). However, genetic separation between nasal and virulent strains had been observed previously (Olvera et al., 2006a and b).

While genotype A was more robust; genotype B had only two strains (HS1075 and FGXBB), which had a different origin but shared a virulent phenotype. These were also characterised as genotype 2 from analysis of the *H. parasuis* population by the MLST method, which were mostly comprised of clinical disease isolates (24/30) (Olvera et al., 2006b). Analysis of further strains is required to determine the robustness of genotype B.

In addition, we analysed the population structure of the *H. parasuis* strains by comparison of the amino acid sequences of the major outer membrane proteins (OMPs) P1, P2 and P5 in an analogous manner to that described recently (Mullins et al., 2009). The derived phylogenetic tree is shown in Fig. 2. There was a clear separation between OMP type A and B lineages and also a unique Chinese OMP profile within the type A lineage, which also contained genetically closer strains 29755, HS1080 and HS80 (Fig. 1). In a comparison between the two polygenetic trees, the low virulent lineage in Fig. 1 was separated as the two OMP lineages in Fig. 2 (line in green), indicating that strains of nasal origin had more heterogeneity in OMP profiles than their genetic profile. Although immune selection may contribute to a variable OMP profile, a relatively convergent Chinese OMP profile demonstrated the Chinese lineage (independent sample *t* test, *P* <0.001).

**Fig. 2.**
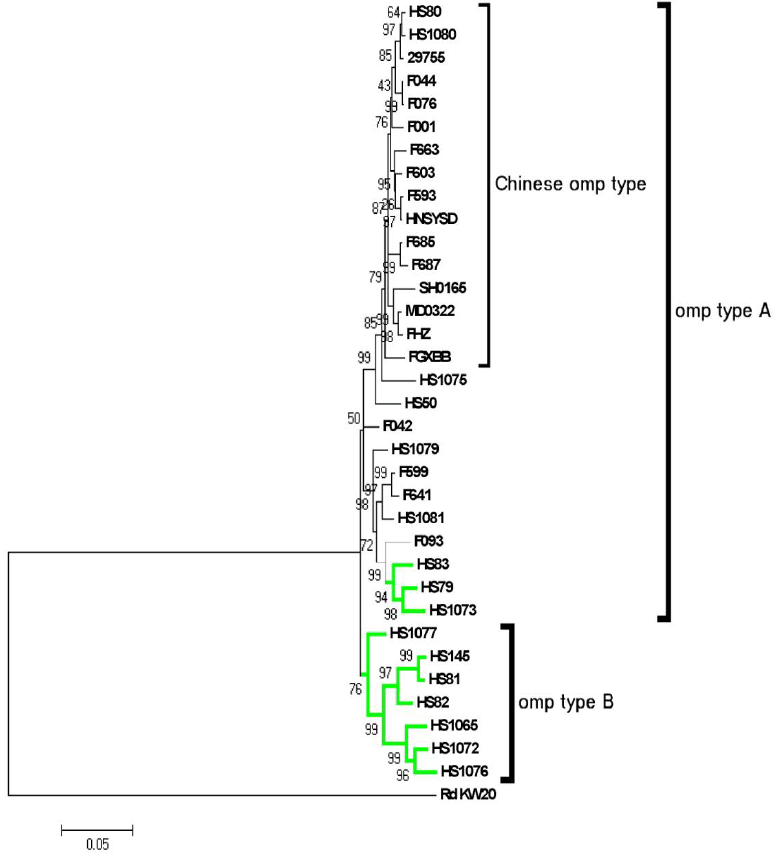
Neighbour-joining (NJ) tree with interior branch test of 2,000 replicates based on concatenated amino acid sequences of the P1, P2 and P5 proteins. Thirty-four *H. parasuis* strains could be divided into two outer membrane protein (OMP) types or profiles (OMP types A and B). OMP type A contained the Chinese OMP type. Green lines indicate nasal isolates. *H. influenzae* strain KW20 was used as the reference for the construction of the phylogenetic tree. The NJ tree was produced by MEGA 4.0. The scale bar is in units of amino acid substitutions per site. Bootstrap values (%) are indicated at the branch nodes of the phylogenetic trees.

Taken together, the data indicate that there are at least two dramatically different genotypes in the *H. parasuis* population. Phylogenetic analysis of both conserved orthologous genes and OMPs indicate there is a unique Chinese lineage, which is associated with a virulent phenotype. Although the first complete genome sequence of Chinese lineage SH0165 gave valuable genetic information, more work needs to be performed targeting which virulence genes result in disease outcome (Yue et al., 2009).

## Acknowledgements

This work was supported by the 973 Program (grant no. 2006CB504404) and Innovation Teams of Ministry of Education (grant no. IRT0726).

## Appendix A. Supplementary material

Supplementary Tables S1 and S2.

---

**Supplementary Table S1.** Strains and background information of Haemophilus parasuis used in the study.

| KRG serovar<sup>a</sup> | Date of receival<sup>b</sup> | Strain | Country of origin<sup>c</sup> | Diagnosis/isolation site | Experimental Infection | Phenotype virulence<sup>d</sup> | Reference |
| --- | --- | --- | --- | --- | --- | --- | --- |
| 1 | 12/86 | HS82 | Japan | Healthy/nose | Glässer's disease | virulent | Amano et al., 1994 |
| 1 | 2/90 | HS145 | Australia | ND | ND | ND | Cai et al., 2005 |
| 2 | 12/86 | HS83 | Japan | Healthy/nose | Healthy | mid-virulent | Nielsen et al., 1993 |
| 3 | 12/86 | HS81 | Japan | Healthy/nose | Healthy | avirulent | Nielsen et al., 1993 |
| 4 | 12/86 | HS79 | Japan | Healthy/nose | Subclinical | mid-virulent | Amano et al., 1994 |
| 5 | 12/86 | HS80 | Japan | Septicaemia/meningitis | Glässer's disease | virulent | Amano et al., 1996 |
| 6 | 9/96 | HS1072 | Switzerland | Healthy/nose | Healthy | avirulent | Nielsen et al., 1993 |
| 7 | 9/96 | HS1073 | Switzerland | Healthy/nose | Healthy | avirulent | Nielsen et al., 1993 |
| 8 | 9/96 | HS1065 | Sweden | ND | Subclinical | mid-virulent | Kielstein et al., 1992 |
| 9 | 11/85 | HS50 | Sweden | ND | Healthy | avirulent | Kielstein et al., 1992 |
| 10 | 9/96 | HS1076 | Germany | Healthy/nose | Glässer's disease | virulent | Kielstein et al., 1992 |
| 11 | 9/96 | HS1077 | Germany | Pneumonia/trachea | Healthy | avirulent | Kielstein et al., 1992 |
| 12 | 9/96 | HS1075 | Germany | Polyserositis/lung | Glässer's disease | virulent | Kielstein et al., 1992 |
| 13 | 9/96 | HS1079 | USA | ND/lung | Glässer's disease | virulent | Kielstein et al., 1992 |
| 14 | 9/96 | HS1080 | USA | ND/joint | Glässer's disease | virulent | Kielstein et al., 1992 |
| 15 | 9/96 | HS1081 | USA | Pneumonia/lung | Polyserositis | virulent | Kielstein et al., 1992 |
| 5 | ND | 29755 | USA | Pneumonia/lung | Glässer's disease | virulent | Nielsen et al., 1993 |
| 5 | 4/01 | SH0165 | Hubei, China | Polyserositis/lung | Glässer's disease | virulent | Cai et al., 2005 |
| 4 | 8/01 | MD0322 | Hebei, China | Polyserositis/lung | Glässer's disease | virulent | Cai et al., 2005 |
| 6 | /03 | F001 | Hubei, China | ND/lung | Glässer's disease | virulent | Cai et al., 2005 |
| 10 | /04 | F042 | Anhui, China | ND/lung | Glässer's disease | virulent | Cai et al., 2005 |
| 7 | /04 | F044 | Henan, China | ND/lung | Glässer's disease | virulent | Cai et al., 2005 |
| 5 | /04 | F076 | Fujian, China | ND/lung | Glässer's disease | virulent | Cai et al., 2005 |
| 2 | /04 | F093 | Hainan, China | ND/lung | Glässer's disease | virulent | Cai et al., 2005 |
| NT | /03 | F593 | Hubei, China | ND/lung | Glässer's disease | virulent | Cai et al., 2005 |
| NT | /04 | F599 | Shandong, China | ND/lung | Glässer's disease | virulent | Cai et al., 2005 |
| NT | /03 | F603 | Henan, China | ND/lung | Glässer's disease | virulent | Cai et al., 2005 |
| NT | /04 | F641 | Henan, China | ND/lung | Glässer's disease | virulent | Cai et al., 2005 |
| NT | /04 | F663 | Shanghai, China | ND/lung | Glässer's disease | virulent | Cai et al., 2005 |
| NT | /04 | F685 | Hunan, China | ND/lung | Glässer's disease | virulent | Cai et al., 2005 |
| NT | /02 | F687 | Hainan, China | ND/lung | Glässer's disease | virulent | Cai et al., 2005 |
| 5 | /04 | FHZ | Zhejiang, China | ND/lung | Glässer's disease | virulent | Cai et al., 2005 |
| 4 | /04 | HNSYSD | Hunan, China | ND/lung | Glässer's disease | virulent | Cai et al., 2005 |
| 5 | /04 | FGXBB | Guangxi, China | ND/lung | Glässer's disease | virulent | Cai et al., 2005 |

<sup>a</sup> There are 15 serovars according to immunodiffusion using heat-stable antigen extracts and NT was defined as the non-typable strains of *H. parasuis* by serotyping method described by Cai. ND indicated the unknown information.
<sup>b</sup> Date expressed in month/year format.
<sup>c</sup> All the 17 Chinese field isolates were collected in 17 distinct farms from 11 provinces representing the Central China (SH0165, MD0322, F001, F044, F593, F603, F641), East China (F042, F076, F599, F663, FHZ), South China (F093, F685, F687, HNSYSD, FGXBB), without disease correlation with available data.
<sup>d</sup> Strains were categorised as the virulent, mid-virulent and avirulent phenotype of *H. parasuis* according to the accompanying reference about experimental infection model. All the 17 Chinese field isolates were provided by the study by Dr. Cai and the detail information about experimental infection with each strains as follows: 9-10 week old weaning piglets (seronegative pigs, 5 for each strain as a group and another 5 for the negative control) were chosen for intraperitoneally challenging with $7 \times 10^9$ CFU, all the infection groups, except for the negative control, showed clinical syndrome as Glässer's disease within a week, and 3~5 piglets in each group died in the following weeks (Cai et al., unpublished data).

**Supplementary Table S2.**
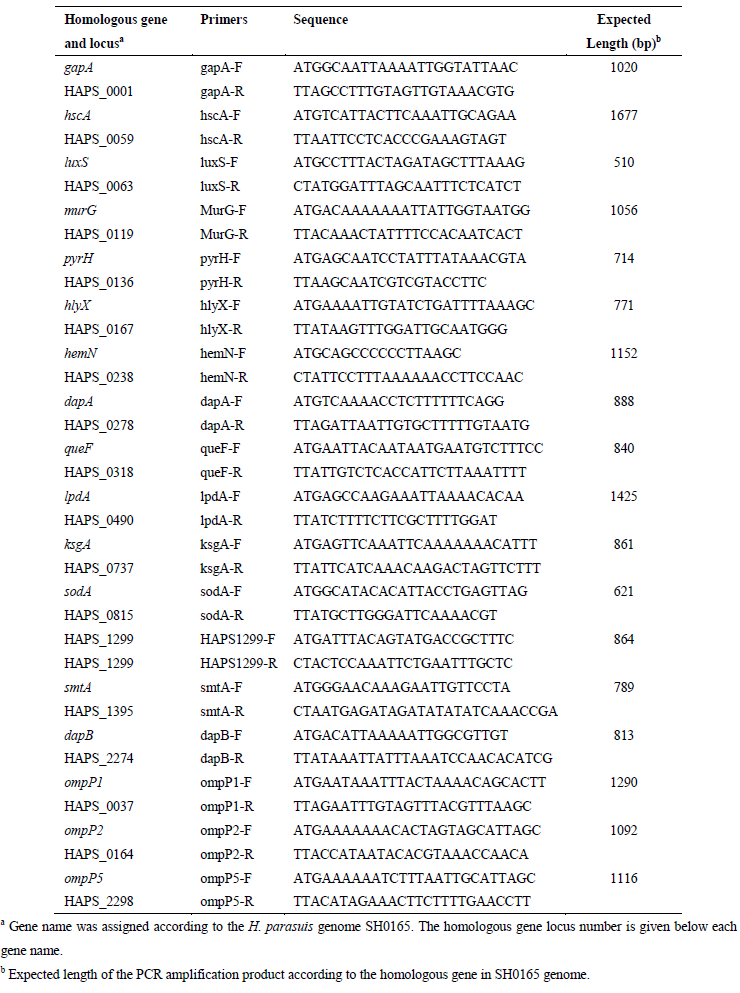
Primers used in this study.

